# Production of brain-derived neurotrophic factor gates plasticity in developing visual cortex

**DOI:** 10.1101/2022.08.04.502872

**Authors:** Megumi Kaneko, Michael P. Stryker

## Abstract

We have previously shown that recovery of visual responses to a deprived eye during the critical period in mouse primary visual cortex (V1) requires both export of mRNA encoding brain derived neurotrophic factor (BDNF) to the dendrites of cortical cells (Kaneko et al., 2012) and phosphorylation of the TrkB receptor for BDNF (Kaneko et al., 2008a). TrkB phosphorylation is not required for the loss of response during monocular deprivation (MD). Studies in vitro show that formation of new connections requires BDNF (Meyer-Franke et al., 1995, 1998; Gottmann et al., 2009). We have now studied the temporal relationship between the production of mature BDNF and the recovery of visual responses. Visual cortical responses to an eye whose vision has been occluded for several days during the critical period and is then re-opened recovers rapidly during binocular vision or more slowly following reverse occlusion, when the previously intact fellow eye is occluded in a model of “patch therapy” for amblyopia. The time to recovery of visual responses differed by 18 hours between these two procedures, but in each the production of mature BDNF preceded the recovery by 6 hours. These findings suggest that a spurt of BDNF secretion stimulates the growth of connections serving the deprived eye to restore visual responses.

**Significance Statement:** By demonstrating that BDNF production precedes rather than merely accompanies the increase in cortical responses, we provide further evidence that BDNF production plays a causal role, thereby gating the plasticity that underlies the recovery of visual cortex from monocular deprivation.

## Introduction

Sensory experience strongly influences maturation and refinement of neuronal connections in the mammalian cortex during postnatal development. In the visual system, depriving visual input by closing one eye (monocular deprivation: MD) for a few days during a critical period of heightened plasticity in early postnatal life leads to a pronounced decrease in the cortical representation of the deprived eye, which is observed both physiologically and anatomically (Wiesel, 1982; Espinosa and Stryker, 2012). A second manifestation of cortical plasticity is the recovery of cortical responsiveness to the closed eye when the vision to the eye is restored. Brain-derived neurotrophic factor (BDNF) has been proposed as a regulator of ocular dominance plasticity (ODP), as visual experience stimulates BDNF expression in the visual cortex (Castren et al., 1992; Lein and Shatz, 2000) and pharmacological interference of signaling mediated by TrkB, the receptor for BDNF, impairs the formation of OD columns (Cabelli et al., 1995, 1997). Using a powerful chemical-genetic approach (Specht et al., 2002; Chen et al., 2005), we previously demonstrated that TrkB signaling during the critical period is required for recovery, but not loss, of cortical responsiveness following MD in mice (Kaneko et al., 2008a). In that study, we chose to examine recovery at 4 days after the deprived eyes were open allowing the animals binocular vision for a similar duration of MD (Kaneko et al., 2008a).

Here we sought to measure the relationship between the production of mature BDNF and the recovery of responses to the deprived contralateral eye in mouse visual cortex. While recovery from the effects of MD requires function of TrkB receptor for BDNF, changes in BDNF production in the visual cortex during the course of recovery have not been examined. As BDNF expression is stimulated by neural activity in the visual cortex (Castren et al., 1992; Bozzi et al., 1995; Schoups et al., 1995; Rossi et al., 1999; Lein and Shatz, 2000; Majdan and Shatz, 2006; Ichisaka et al., 2003; Pattabiraman et al., 2005; Karpova et al., 2010; Schwartz et al., 2011), up-regulation of BDNF is expected during recovery. It seemed therefore from the literature that BDNF production might follow and be a result of the increased cortical responses to the recovering eye’s pathways. What we found instead is that an increase in BDNF production precedes the increase in the efficacy of deprived-eye pathways by at least the shortest time interval that we measured, 6 hours. BDNF production preceded the recovery of responses by a similar amount in two different circumstances—with rapid recovery during binocular vision as well as with much slower recovery following reverse occlusion. These findings are consistent with a causal role for BDNF in strengthening excitatory pathways in the cortex.

## Results

### Recovery of deprivation-induced depression in cortical responses is faster during binocular vision than during reverse occlusion

We examined recovery using chronic optical imaging of intrinsic signals in primary visual cortex (V1) during the developmental critical period to measure the magnitude of cortical responses to visual stimuli repeatedly in the same animals before eyelid suture, after 5 days of MD, and after various durations of either binocular vision (BV) or reverse occlusion (RO) (Fig. 1A). Note that different groups of mice were used for each time point because the necessity for short durations of recovery to compare RO and BV precluded repeated exposure of animals to anesthetics. Figure 1B shows examples of intrinsic signal images after MD followed by 2 days of BV.

**Figure 1.**
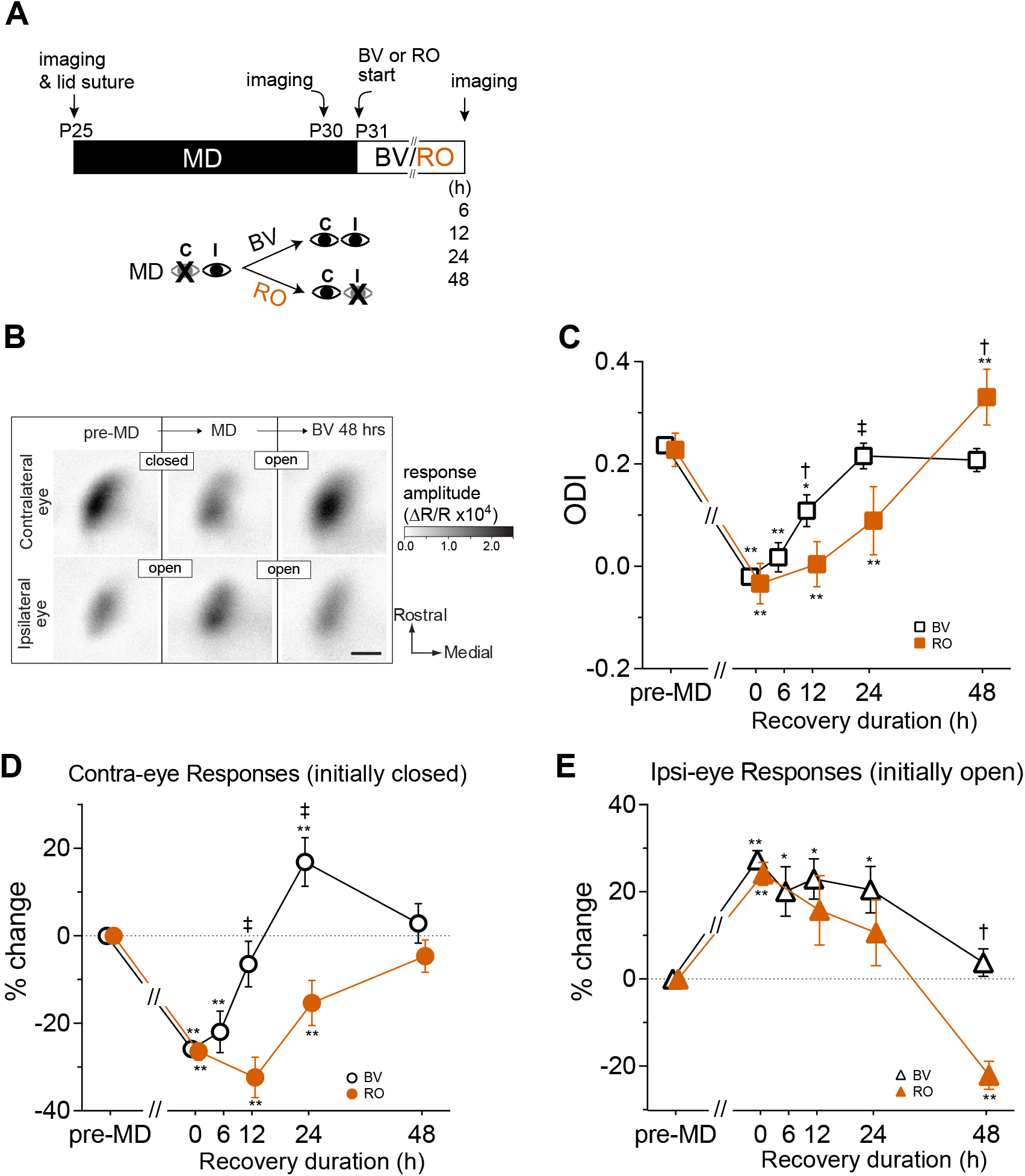
Faster recovery of deprived eye responses by binocular vision than by reverse occlusion A. Experimental schedule for repeated optical imaging of intrinsic signals. MD: monocular deprivation of contralateral eye. BV: binocular vision, open symbols. RO: reverse occlusion, filled symbols. B. Example of changes in response magnitude in an animal undergoing MD followed by recovery by BV or RO. The gray scale represents the response magnitude as a fractional change in reflectance. Scale bar = 0.5 mm for all panels. C. Shift in ocular dominance during recovery calculated from data in D and E. D. Changes in the response magnitude of the contralateral eye (closed → open) eye during recovery. E. Changes in the response magnitude of the ipsilateral eye (open → open for BV, open → closed for RO) during recovery. Data in D and E are presented as % change from pre-MD baseline. All graphs show mean ± s.e.m. Sample size: BV-6h (4), BV-12h (5), BV-24h (5), BV-48h (5), RO-12h (5), RO-24h (5), RO-48h (6). *P<0.05, **P<0.01 vs. pre-MD baseline, repeated measure ANOVA followed by multiple comparisons with Bonferroni’s correction. †P0.05, ‡P<0.01 between BV and RO, one-way ANOVA followed by multiple comparisons with Bonferroni’s correction.

Previous studies showed recovery of ocular dominance by 4 days of binocular vision (Kaneko et al., 2008a) or 4 days of reverse occlusion (Gordon and Stryker, 1996) following 4 – 6 days of MD during the critical period. Therefore, we examined effects of BV or RO for duration of 2 days or shorter (Fig. 1A). Recovery of ODI occurred faster with BV than with RO. It was already evident at BV-12 h (MD: −0.020 ± 0.036, BV-12h: 0.109 ± 0.062), was complete by 24 h (0.216 ± 0.051), and no further change at 48 h (0.208 ± 0.045) (Fig. 1C). In contrast, in mice with RO, a partial recovery to the level similar to BV-12h was detected only after 24 h (0.089 ± 0.066) (Fig. 1C). Complete recovery of ODI under RO seemed to take place sometime between 24h and 48h (Fig. 1C).

Recovery in ODI at BV-48h was a result of both full restoration of open-eye responses and returning of closed-eye responses to the baseline level (Fig. 1D, E). At BV-24h and BV-12h, changes in the magnitude of closed-eye responses were different from those to the open-eye. At BV-12h, closed-eye responses were restored to −6.5 ± 10.4% of baseline (P>0.05, Fig. 1D) but open-eye responses were still at an elevated level similar to those observed immediately after MD (BV-12h: 22.9 ± 9.3%, MD: 27.3 ± 8.6% of baseline, Fig. 1E). At BV-24h, closed-eye responses were mildly but significantly increased beyond the baseline (16.9 ± 11.1%, P<0.05, Fig. 1D) while open eye responses were still at an elevated level (20.5 ± 10.6% of baseline, P>0.05 vs. MD, Fig. 1E), resulting in an apparently normal ODI.

Delayed recovery of ODI with RO was a result of a slower increase in previously closed-eye responses, which seemed to take 48 h for full recovery to the baseline (Fig. 1D). At this time point, responses to the previously open, newly closed, eye were depressed (−22.1 ± 7.2% of baseline, P<0.05, Fig. 1E), resulting in significant OD shift toward the previously closed eye (RO-48h: 0.331 ± 0.055, baseline: 0.218 ± 0.032, P<0.05, Fig. 1C). These results showed that responses to the previously closed eye recovered faster when both eyes were open than when the occlusion was reversed following MD. In addition, recovery of responses to the previously closed eye during BV has a distinct time course from that of previously open eye responses.

### Requirement of TrkB kinase function for rapid recovery

A Phenylalanine-to-alanine substitution (TrkB^F616A^) within the ATP binding pocket of the kinase renders the receptor sensitive to specific inhibition by small molecule 1NMPP1, a derivative of the general kinase inhibitor PP1, allowing for specific and temporally controlled inactivation of TrkB (Chen et al., 2005). Using these transgenic mice, we have shown that tyrosine kinase activity of the TrkB neurotrophin receptor is required for full recovery from effects of MD during 4 days of BV (Kaneko et al., 2008a). Because we found that recovery occurred much faster as described above, we examined effects of TrkB inactivation during 2 days of BV or RO.

TrkB^F616A^ homozygous mice were given 1NMPP1 or vehicle solution via osmotic minipump throughout the duration of BV (12, 24, or 48 h) or RO (48h) (Fig. 2A). Consistent with our previous result for 4d-BV, TrkB inactivation greatly reduced the restoration of cortical responses to the previously closed eye during 48 h of BV (vehicle: 2.9 ± 5.9%, 1NMPP1: −16.7 ± 8.4 %, P<0.05) (Fig. 2B). However, it showed a modest and transient increase between 0 h and 24 h (0h: −29.4 ± 5.8 %, 24h: −10.2 ± 9.3 %, P<0.05). Responses to the previously open eye in 1NM-PP1-treated mice did not change during the whole period of 48 h BV, that is, stayed at the same elevated level of MD (Fig. 2C). As a result, the ocular dominance was partially restored at 24 h (0h: −0.03 ± 0.06, 24h: 0.13 ± 0.05, P<0.05) but it was transient and fell back to a level similar to that of MD at 48 h (0.07 ± 0.05) (Fig. 2D).

**Figure 2.**
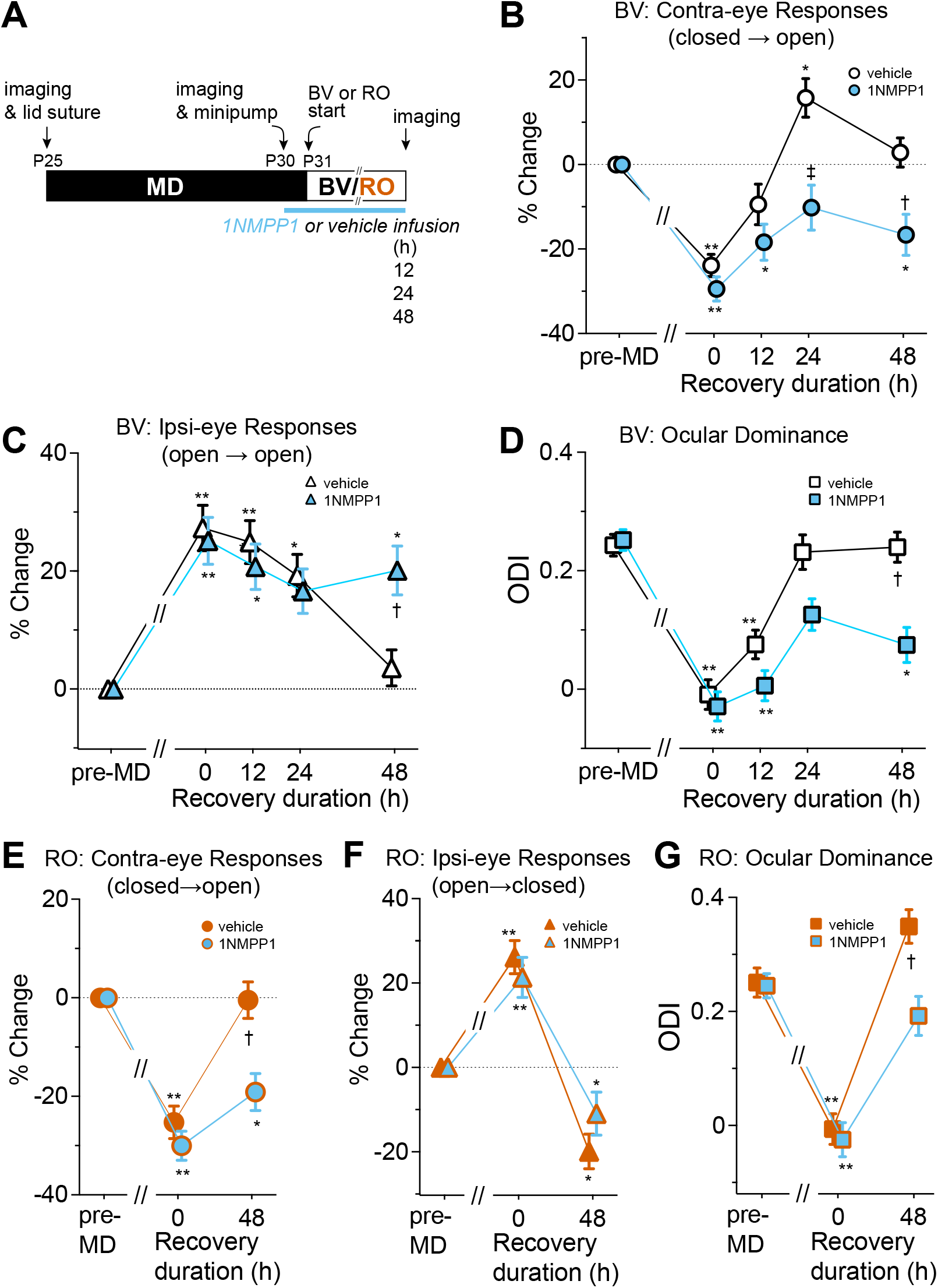
Effects of blocking TrkB receptor function on the time course of recovery. A. Experimental schedule. B - D. Recovery by binocular vision (BV) in TrkB^F616A^ mice treated with vehicle or 1NMPP1 infusion. %Change in magnitude of contralateral eye (closed → open) responses (B) and of ipsilateral eye (open → open) responses (C), and ocular dominance index (D). Sample size for vehicle-treated animals: 12 h (4), 24 h (5), 48 h (5); for 1NMPP1-treated animals: 12 h (4), 24 h (5), 48 h (5). E - G. Recovery by reverse occlusion in TrkB^F616A^ mice treated with vehicle or 1NMPP1 infusion, %change in the response magnitude to the contralateral eye (closed → open) (E) and to the ipsilateral eye (open → closed) (F), and ocular dominance index (G). *P<0.05, **P<0.01 vs. pre-MD baseline, repeated measure ANOVA followed by multiple comparisons with Bonferroni’s correction. †P0.05, ‡P<0.01 between vehicle and 1NMPP1, one-way ANOVA followed by multiple comparisons with Bonferroni’s correction.

Similar to its effect during BV, TrkB inactivation inhibited recovery of responses through the previously closed eye during RO (vehicle: −0.47 ± 7.4 %, 1NM-PP1: −19.0 ± 7.5 %, P<0.01) (Fig. 2E). In contrast, it did not inhibit the decrease in responses to the previously open, newly closed eye (vehicle: −19.9 ± 8.2 %, 1NMPP1: −11.0 ± 10.2 %, P>0.05) (Fig. 2F). Consequently, ocular dominance in TrkB inactivation group was shifted toward the previously closed eye during RO as much as the baseline pre-deprivation level (1NMPP1: baseline: 0.25 ± 0.04, RO: 0.19 ± 0.07, P>0.05), but was significantly less so compared to the control group (vehicle: baseline 0.25 ± 0.05, RO: 0.35 ± 0.06) (Fig. 2G). This lack of effect for TrkB inactivation on depression of responses to the newly closed eye is consistent with our previous observation that it was not required for decrease in closed-eye responses during MD (Kaneko et al., 2008a).

### Upregulation of BDNF protein during physiological recovery

Neural activity-dependent expression of BDNF has been extensively documented (Lessmann et al., 2003; Lu, 2003; Tongiorgi, 2008; Park and Poo, 2013). Its expression in the visual cortex is modulated by visual experiences and is down-regulated during MD (Castren et al., 1992; Bozzi et al., 1995; Rossi et al., 1999; Lein and Shatz, 2000; Ichisaka et al., 2003; Majdan and Shatz, 2006). We examined the changes in BDNF protein level within the binocular area of V1 during BV or RO (Fig. 3A). We used Western blot analysis (Fig. 3B) because it allows distinction between mature BDNF and pro-BDNF; the former, a proteolytic product of the latter, is the high-affinity ligand for TrkB (Reichardt, 2006).

**Figure 3.**
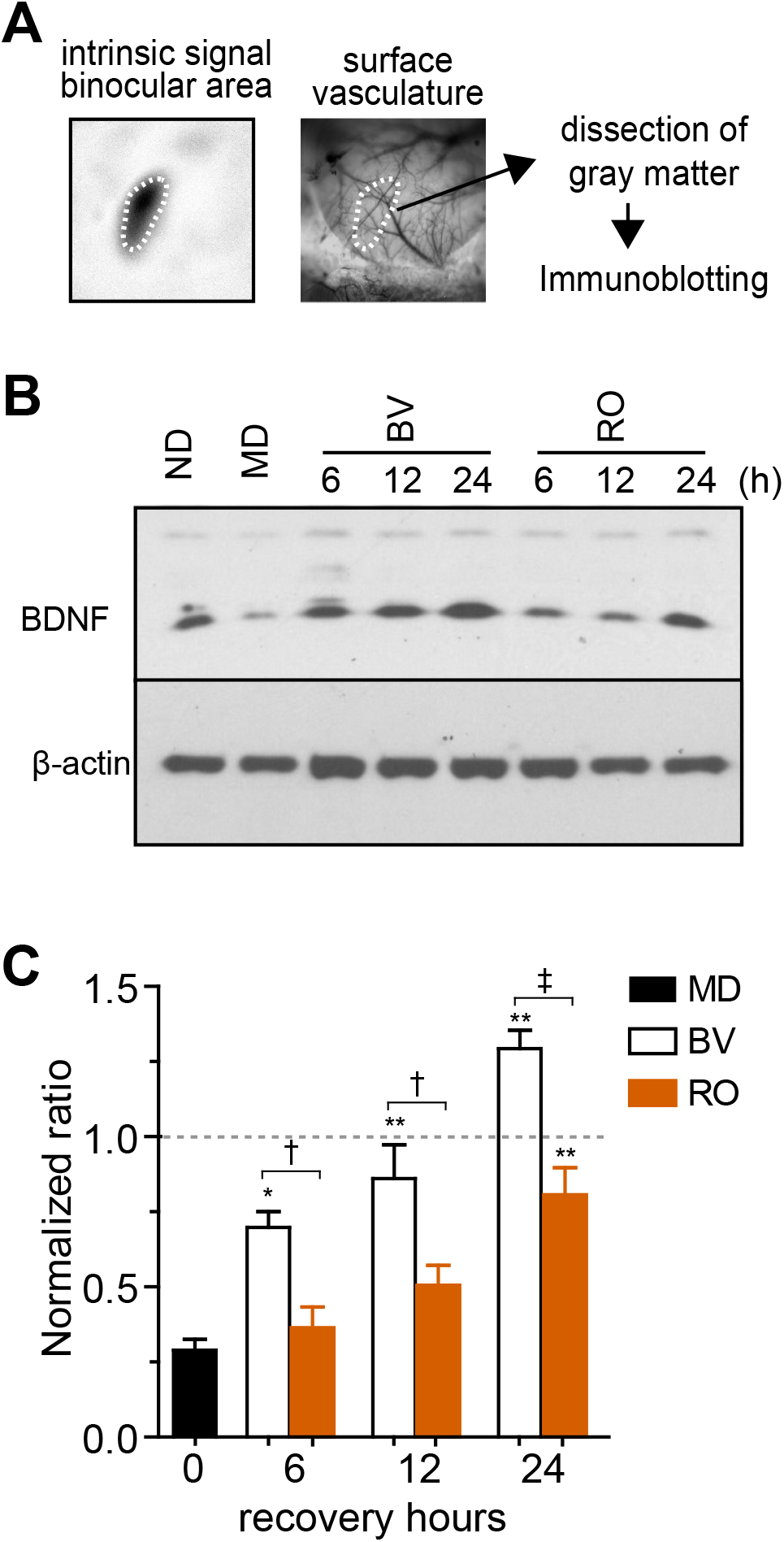
Rapid upregulation of BDNF in the primary visual cortex during recovery by binocular vision. A. Dissection of the binocular visual cortex that has been localized using the surface vascular pattern and the intrinsic signal image. B. An example of the immunoblot for BDNF. Beta-actin was used as a loading control. C. Quantification of density of the immuno-blot. The magnitude of the BDNF signal was first normalized with the corresponding beta-actin signal and then normalized with the magnitude of no deprivation (ND) control in the same experiment. Data are presented as mean ± s.e.m. Sample size is 5 pairs for each condition. MD: monocular deprivation, BV: binocular vision following 6 days of MD, RO: reverse occlusion following 6 days of MD. *P<0.05 and **P<0.01 vs. ND; †P0.05 and ‡P<0.01 between BV and RO; all statistical analyses were done by one-way ANOVA followed by multiple comparisons with Bonferroni’s correction.

Consistent with previous studies, mature BDNF (mBDNF) protein level was significantly downregulated in the binocular area immediately after 5 days of MD (29.0 ± 5.1%, P<0.05 vs. ND). Upregulation of mBDNF during the recovery period occurred faster with BV than with RO (Fig. 3C). There was a partial but significant increase from MD level already at BV-6h (69.9 ± 5.2%, P<0.05 vs. MD) and it reached almost to the ND level at BV-12h (86.1 ±11.3%, P>0.05 vs. ND). In addition, mBDNF level at BV-24h increased slightly but significantly beyond the ND level (129.3 ± 6.1%, P<0.05 vs. ND). Under RO, mBDNF did not return to the normal level until 24h (80.1 ± 9.1%, P<0.01 vs. MD) with no significant increase detected at 6h or 12h (6h: 36.4 ± 6.9%, 12h: 50.6 ± 9.5%, both P>0.05 vs. MD).

How do changes in the BDNF level relate to functional recovery? At BV-6h, mature BDNF was already significantly increased from that after MD (Fig. 3C), whereas the previously closed-eye responses were on average still at MD level (Fig. 1B). Also, at RO-24h significant increase was detected in mature BDNF (Fig. 2C) but not in previously closed eye responses (Fig. 1B). Thus, during both BV and RO, the increase in mature BDNF protein level preceded functional recovery of the responses to the previously closed-eye.

### Homeostatic synaptic scaling during MD contributes to rapid recovery by BV

When overall cortical activity is reduced, the strength of excitatory synapses increases, a process referred to as homeostatic synaptic scaling. This process is mediated at least in large part by signaling through tumor necrosis factor alpha (TNFα) (Kaneko et al., 2008b; Stellwagen and Malenka 2006). During MD, the initial decrease in closed-eye responses lowers overall cortical activity, thereby engaging TNFα-dependent homeostatic scaling to increase responses to the open eye (Kaneko et al., 2008b). Homeostatic scaling also works on closed-eye responses to increase it to a lesser degree. We tested whether such a homeostatic increase in responsiveness through TNFα signaling contributes to the faster recovery of the previously closed eye responses and changes in ocular dominance.

Consistent with our previous study (Kaneko et al, 2008b), partial blockade of TNFα signaling by cortical infusion of soluble TNF receptors (sTNFR) during 5 days of MD almost completely suppressed the normal increase in open-eye responses and caused a slightly larger decrease in closed-eye responses, resulting in a much smaller ODI shift compared to vehicle control (points at MD in Fig. 4A - D). In these animals, 24 h of BV produced no significant change either in responses to the previously closed eye or in ocular dominance. Instead it took 2 days for significant changes to occur (Fig. 4B, D). This effect of blocking TNFα signaling was to reduce the strength of the synapses that would otherwise have increased in strength, as they did with vehicle infusion. The synapses left less powerful by the blockade of TNFα signaling would presumably produce less of the cortical activity that stimulates secretion of mature BDNF.

**Figure 4.**
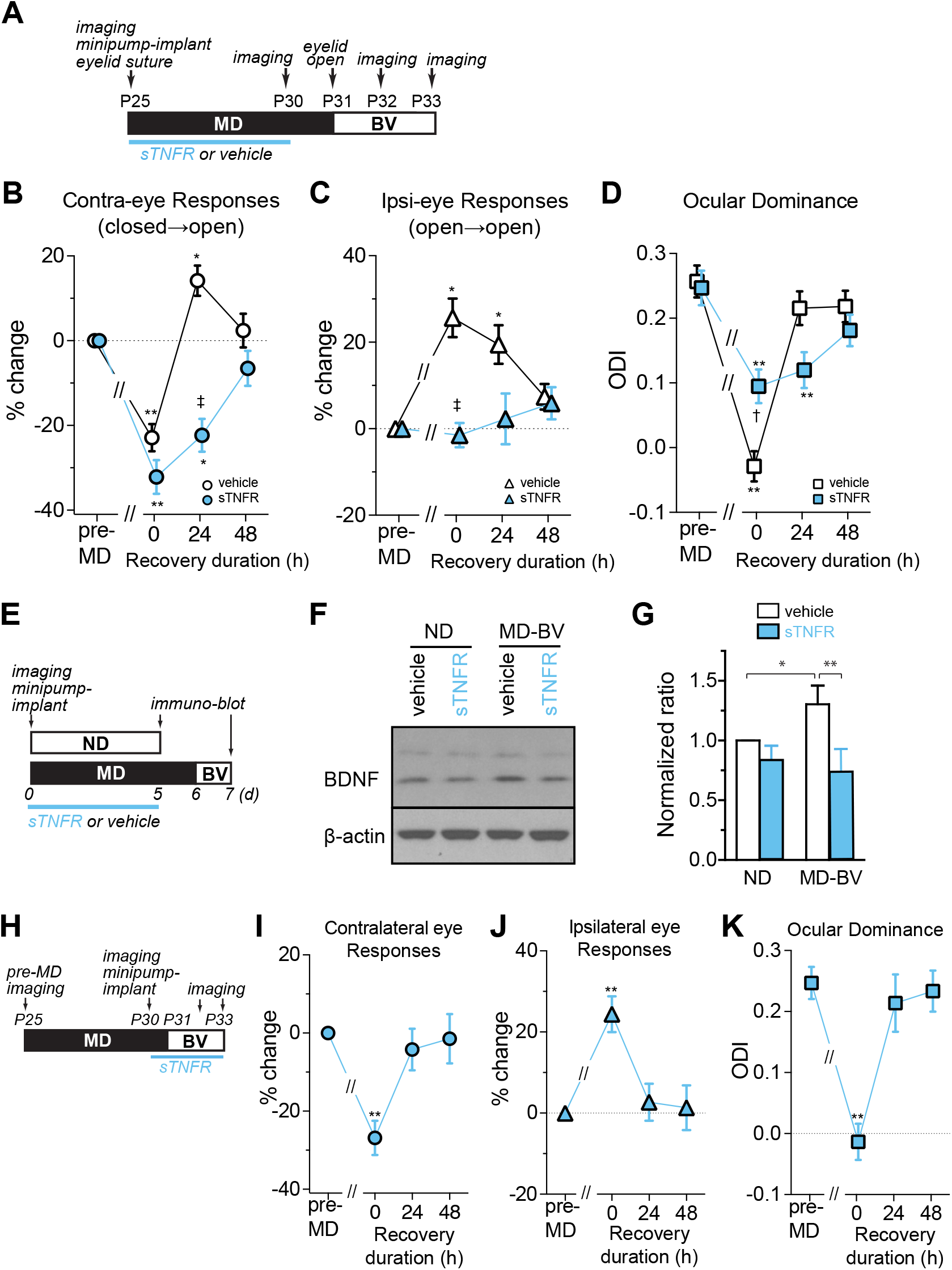
TNFα signaling during MD contributes to rapid functional recovery and BDNF upregulation. A. Experimental schedule. B – D. Changes in responses to the contralateral eye (closed → open, B); to the ipsilateral eye (open → open, C); and in ocular dominance (D) during recovery by binocular vision (BV) following 6 days of MD. Dotted lines in B and C represent baseline level. The response magnitude is shown as % change from pre-MD baseline. Data are presented as mean ± s.e.m. Sample size: vehicle (5, open symbols in black), soluble TNF receptor (sTNFR) (5, symbols filled with blue). *P<0.05 and **P<0.01 vs. pre-MD baseline, repeated measure ANOVA followed by multiple comparisons with Bonferroni’s correction. †P0.05 and ‡P<0.01 between vehicle and sTNFR groups, one-way ANOVA followed by multiple comparisons with Bonferroni’s correction. E – G. Effects of sTNFR infusion on BDNF level. (E) Experimental schedule. (F) An example of the BDNF immunoblot. (G) Quantification of the BDNF level. Signal strength for BDNF was first normalized to the corresponding beta-actin signal and then normalized to that of no-deprivation vehicle control in each blot. *P<0.05 and **P<0.01, one-way ANOVA followed by multiple comparisons with Bonferroni’s correction. H - K. Effects of sTNFR infusion during recovery by BV sTNFR infusion was started immediately after post-MD imaging, followed 1 day later by the start of binocular vision (H). Changes in responses to the contralateral eye (closed → open, I); to the ipsilateral eye (open → open, J), and in ocular dominance (K). Data are shown as mean ± s.e.m. **P<0.01 vs. pre-MD baseline, one-way ANOVA followed by multiple comparisons with Bonferroni’s correction.

Indeed, the delayed recovery of function under TNFα blockade was accompanied by reduced BDNF upregulation. In mice treated with sTNFR infusion, the levels of mature BDNF after 24 h of BV were significantly lower than vehicle control animals after 24 h of BV and were similar to those in non-deprived mice infused with vehicle solution (BV-sTNFR mBDNF was 73 ± 19 % of ND-veh). In contrast, in animals treated with vehicle solution, mature BDNF levels after 24h of BV were significantly increased to 130 % of the levels in non-deprived vehicle control animals (Fig. 4F, G), similar to the increased level we had measured in animals after 24h of BV with no cortical infusion, as shown in Figure 3.

A potential complication is that sTNFR might have not cleared completely when BV was started so that residual sTNFR might have affected recovery. However, this possibility is unlikely because infusion of sTNFR starting 1 day before and throughout the duration of 48 h of BV (Fig. 4H) had no detectable effect on recovery of responses, which was similar to that in control BV animals with no infusion (Fig. 4I, J, K). Thus, increase in synaptic strength during deprivation, mediated by TNFα signaling, is necessary for both the spurt of BDNF secretion and for the rapid recovery of cortical responses during the first 24 h of restored binocular vision.

This finding provides further support for the coupling between the section of mature BDNF and recovery of cortical responses to the deprived eye.

## Discussion

In nearly pure neuronal cultures studied *in vitro*, the formation of new synapses and the activity-dependent strengthening of existing ones requires signaling by brain derived neurotrophic factor (BDNF) on its principal receptor, TrkB (Meyer-Franke et al., 1995, 1998; Gottmann et al., 2009; Park and Poo, 2013; Kowiański et al., 2018; Horch and Katz, 2002; Tanaka et al., 2008; Harward et al., 2016). *In vivo,* neural activity has long been known to stimulate the production of mature BDNF (Castren et al., 1992; Bozzi et al., 1995; Schoups et al., 1995; Rossi et al., 1999; Lein and Shatz, 2000; Ichisaka et al., 2003; Pattabiraman et al., 2005; Majdan and Shatz, 2006; Karpova et al., 2010; Schwartz et al., 2011; Miyasaka et al., 2021). To determine whether the recovery of responses after the cessation of MD to the re-opened, formerly deprived, eye requires BDNF-TrkB signaling, and is therefore likely to rely on the strengthening of existing synaptic connections and the formation of new ones, we used a chemical-genetic approach (Kaneko et al., 2008a). We studied recovery in mice in which TrkB receptors had been engineered to be susceptible to a small-molecule “Shokat” inhibitor (Specht et al., 2002; Chen et al., 2005). Application of the inhibitor in these mice completely blocked the recovery of deprived-eye responses, as well as the apparently homeostatic reduction in the responses to the open eye. These findings provided strong evidence that BDNF secretion mediated an essential step in the recovery of deprived-eye responses.

To test this hypothesis further, here we examined the time course of the appearance of mature BDNF in relation to the recovery of deprived-eye responses after the re-opening of the deprived eye. If BDNF plays a causal role in recovery, then it should appear before responses recover when BDNF production and responses are examined with sufficient time resolution. Here we studied recovery of deprived-eye responses under two different conditions: one in which the deprived eye was opened while the fellow eye remained open; and a second, referred to a “reverse occlusion”, in which the fellow eye was deprived of vision by lid suture at the time the initially deprived eye was opened. The former condition produces rapid recovery, in less than 24 hours; recovery in the latter condition is much slower, presumably because neither eye drives the cortex well immediately after reverse suture. In both cases, the appearance of mature BDNF as measured with Western blots preceded that of recovery of deprived-eye responses by 6 hours or more, consistent with a role in promoting the growth of synapses serving that eye’s pathways. In the case of the more rapid recovery, BDNF levels and responses actually overshot before returning to normal levels.

### A model of the role of BDNF signaling in monocular deprivation and recovery

These and other findings support the following model of the events responsible for monocular deprivation and recovery. Occluding the vision of the contralateral eye, which is the dominant eye in the mouse, dramatically reduces cortical activity and leads to the loss of synapses serving that eye (Sun et al., 2019). The reduction in cortical activity stimulates homeostatic synaptic scaling, which makes the remaining synapses stronger through a process mediated by TNFα signaling, leading to some recovery of cortical activity, a large increase in responses to the intact fellow eye, and a smaller increase in the responses to the deprived eye. When the deprived eye is re-opened, while its pathway’s synapses are stronger than they would otherwise be by virtue of synaptic scaling, they are also fewer than they once were. However, opening that eye, and leaving the fellow eye open as well, immediately increases cortical activity to a great extent, leading to a spurt of mature BDNF secretion. The BDNF acts on TrkB receptors to strengthen existing synapses serving the deprived eye and to promote the formation of new ones in the newly active pathways serving the formerly deprived eye.

In contrast, during reverse occlusion when the fellow (ipsilateral) eye is closed at the time that the deprived (contralateral) eye is re-opened, cortical activity is not increased; at the end of 4 days of MD in the mouse, the two eyes drive the cortex more or less equally well, so that closing one when the other is opened does not change the total excitation of the cortex. In this case, the secretion of mature BDNF is not stimulated and remains at its constitutive level. This level of BDNF permits the strengthening and addition of synapses in the pathway serving the initially deprived (contralateral) to proceed only slowly, leading to a protracted recovery of its responses. Note that this process has positive feedback: as the deprived eye synapses strengthen and new ones are added, they activate the cortex more powerfully so that by 24 hrs of monocular recovery BDNF secretion reaches a level similar to that after 6 hrs of binocular recovery.

This model, which describes the results in Figure 1, is supported by the results in Figure 2, in which BDNF signaling is attenuated using the chemical-genetic inhibitor of its TrkB receptor. Recovery of deprived (contralateral) eye responses during binocular vision is dramatically slowed, and open (ipsilateral) eye responses, which had increased during MD, remain elevated.

The rapid production of mature BDNF during binocular vision, shown in Figure 3 provides further support for this model of deprivation and recovery, as does its delayed appearance during reverse occlusion.

The reduction of homeostatic synaptic scaling illustrated in Figure 4 provides additional support for the model. Blocking synaptic scaling with sTNFR reduces both the increase in cortical activity triggered by binocular vision immediately upon reopening the deprive eye and the consequent increase in the production of mature BDNF. The increase in responses to the deprived eye are delayed, presumably until constitutive BDNF production suffices for enough synapses serving the deprived eye to be strengthened or formed.

Of course, BDNF production is nowhere near that whole story. The role posited here for BDNF signaling does not specify which synapses will be strengthened. In this model BDNF is thought merely to be permissive for the strengthening of synapses and the formation of new ones. The specification of which synapses are to be strengthened or added is presumably a result of normal Hebbian plasticity, with its conventional limits that keep it from running away completely.

The model as presented is abstract. Of course the eyes do not project to the cortex, and within the cortex evidence indicates that plasticity is much more rapid in the upper layers than in the layers that receive the largest input from the lateral geniculate nucleus (Trachtenberg et al., 2000). So when we speak of strengthening the deprived eye pathways, those pathways span at least two synapses between the eyes and the site of the most rapid plasticity in the visual cortex. The abstract nature of this account is made further clear by the ongoing turnover of synapses in the primary visual cortex during the critical period, which encompasses the entire time course of the current study. The net effect of occluding the contralateral eye is a reduction in the number of synaptic boutons in layer 2/3 (Sun et al., 2019, Figure 3D). Subsequent binocular vision then restores the number of boutons to control levels.

With these limitations, however, the model of deprivation and recovery and the proposed role for BDNF signaling in that process has strong experimental support. The most important finding of this study is that the BDNF production precedes the increase in the efficacy of deprived-eye pathways and therefore may play a causal role.

### More rapid recovery with binocular vision than with reverse occlusion

More than a decade ago, we showed that visual cortical responses to a contralateral eye deprived of vision for several days recover much more rapidly with binocular vision than with reverse occlusion in the mouse (Kaneko et al., 2010). This finding was initially surprising in light of the evidence from monkey studies, where binocular vision was seen to be ineffective, and from the clinical experience with amblyopic patients, where “patch therapy” or some form of reverse occlusion is the standard and generally effective therapy (LeVay et al., 1980; Meier and Tarczy-Hornoch, 2022; Repka, 2022). However, a number of additional reports in various experimental animals confirmed that binocular vision could be effective in restoring visual responses after monocular deprivation and sometimes more effective than reverse occlusion (Mitchell et al., 2001; Kind et al., 2002; Faulkner et al., 2006; Mansouri et al., 2014; Mitchell and Duffy, 2014; Hess and Thompson, 2015; Murphy et al., 2015; Jin et al., 2022).

The reason for the difference between rodents on the one hand and human and non-human primates on the other need not be the underlying cellular neurobiology. Instead the difference may be due to the dramatic imbalance in the mouse between the contralateral and ipsilateral visual pathways. In the mouse contralateral-eye pathway provides 4-10 times as many inputs to even the most binocular portion of the visual cortex as do ipsilateral-eye pathways, and the contralateral eye is much more effective than the ipsilateral eye in driving most cortical neurons. In the mouse, several days of contralateral-eye deprivation results in a visual cortex in which the two eyes are more or less equally effective. In humans and macaque monkeys, the strengths and numbers of inputs from the two eyes are similar to each other, and a similar period of monocular deprivation leaves both the ipsilateral or contralateral visual cortex nearly unresponsive to the deprived eye, leaving responses to be driven almost solely by the open eye. In the mouse, binocular vision provides more or less twice the cortical activity and BDNF production as reverse occlusion immediately upon eye opening. Because visual responses to the deprived eye are almost absent in monkeys and humans, binocular vision immediately upon eye opening stimulates about the same amount of cortical activity, and presumably BDNF production, as does reverse occlusion.

While the present mouse findings on the superiority of binocular vision over reverse occlusion may have little direct relevance to the treatment of human amblyopia, the present finding that BDNF production precedes and appears to be casual for the strengthening of a cortical pathway may indeed have great import for therapy. These findings raise the possibility that application of an artificial activator of the TrkB receptor might prove a powerful adjunct to physical or cognitive therapy for brain injuries and disorders (reviewed in Liu et al., 2016; Pankiewicz et al., 2021; Tuszynski 2022).

## Methods

### Animals

All procedures were approved by the Institutional Animal Care and Use Committee of University of California San Francisco. C57/BL6 wild type mice were purchased from Jackson Laboratories and bred in the UCSF animal care facility. TrkB^F616A^ mutant mice were as described (Chen et al., 2005; Kaneko et al., 2008a). Animals were kept in the standard housing condition with a 12h-light/12-dark cycle and a free access to food and water.

Monocular deprivation was performed by suturing the lid of the right eye (contralateral to the imaged hemisphere) at P25 as described (Kaneko et al., 2008b). To examine recovery from MD, mice were imaged immediately before MD, after 5 days of MD, and at one of the time points (6, 12, 24, or 48 h) after restoring vision to the deprived right eye (following 6 days of MD) either by simply removing the suture (binocular vision, BV), or by reversing the lid suture (reverse occlusion, RO).

Continuous infusion of 1NMPP1 or vehicle solution was made into TrkB^F616A^ homozygous mice as described (Kaneko et al., 2008a). Briefly, immediately after optical imaging of intrinsic signals on MD-5d, an osmotic minipump (Alzet model 2001, Cupertino, CA) containing 1NMPP1 (0.25 nmol/ g body weight/ h) or vehicle solution was implanted subcutaneously. In addition, two intraperitoneal injections (16.6 ng/ g body weight) were made during the first 24 h of infusion to facilitate TrkB inactivation. Intracortical infusion of sTNFR or vehicle solution was performed using osmotic minipump (model 1002, Alzet) as described (Kaneko et al., 2008b) during MD or BV Briefly, a cannula was implanted into the medial edge of V1, immediately after intrinsic signal imaging of pre-MD baseline responses (for infusion during MD) or of post-MD responses (for infusion during BV). The cannula was connected to an osmotic minipump filled either with vehicle solution (PBS containing 0.1% BSA as a carrier) or 35 μg/ml of soluble TNF receptor-1 (sTNFR1, R&D Systems Inc. Minneapolis, MN).

### Repeated optical imaging of intrinsic signals

Repeated optical imaging of intrinsic signals and quantification of ocular dominance were performed as described (Kaneko et al., 2008b). Briefly, during recording mice were anesthetized with 0.7% isoflurane in oxygen applied via a home-made nose mask, supplemented with a single intramuscular injection of 20 - 25 μg chlorprothixene. Images were recorded transcranially; the scalp was sutured closed at the end of each session and re-opened at the same location in subsequent sessions. Intrinsic signal images were obtained with a Dalsa 1M30 CCD camera (Dalsa, Waterloo, Canada) with a 135 x 50 mm tandem lens (Nikon Inc., Melville, NY) and red interference filter (610 ± 10 nm). Frames were acquired at a rate of 30 fps, temporally binned by 4 frames, and stored as 512 x 512 pixel images after binning the 1024 x 1024 camera pixels by 2 x 2 pixels spatially. The visual stimulus for recording the binocular zone, presented on a 40 x 30 cm monitor placed 25 cm in front of the mouse, consisted of 2°-wide bars, which were presented between −5° and 15° on the stimulus monitor (0° = center of the monitor aligned to center of the mouse) and moved continuously and periodically upward or downward at a speed of 10°/sec. The phase and amplitude of cortical responses at the stimulus frequency were extracted by Fourier analysis as described (Kalatsky and Stryker, 2003). Response amplitude was an average of at least 4 measurements. Ocular dominance index was computed as described (Cang et al., 2005). Briefly, the binocularly responsive region of interest (ROI) was chosen based on the ipsilateral eye response map after smoothing by low-pass filtering using a uniform kernel of 5 × 5 pixels and thresholding at 40 % of peak response amplitude. We then computed OD score, (C–I)/(C+I), for each pixel in this ROI, where C and I represent the magnitude of response to contralateral and ipsilateral eye stimulation, followed by calculation of the ocular dominance index (ODI) as the average of OD score for all responsive pixels.

### Western blot analysis

The location of the binocular area in the primary visual cortex was identified based on the intrinsic signal map and the photograph of the surface blood vessels that were obtained before starting MD Fig. 3A). The start time of recovery by BV or RO was adjusted so that all experimental groups were sampled at a similar time (early afternoon) to avoid possible circadian influences. The gray matter of binocular V1 contralateral to the initially-closed eye was dissected out under isoflurane anesthesia, frozen in liquid nitrogen, and stored at −80°C. The samples from 2 animals of the same condition were pooled so that one lane in the blot contains two binocular V1. The tissues were homogenized with 5 volume of lysis buffer containing 50mM Tris-HCl buffer (pH 7.5), 150mM NaCl, 1mM EDTA 1% Triton-X 100, 1% Na deoxycholate, 1% SDS, and protease inhibitor cocktail (Sigma, St. Louis), followed by centrifugation and collection of supernatants. Approximately 20 μg of total proteins in tissue lysates were separated on electrophoresis using 4 - 12% gradient SDS-polyacrylamide gels (BioRad) and transferred to the PVDF membrane (BioRad). After blocked with 5% skim milk in 0.05% Tween-20/Trisbuffered saline (TBST), the membrane was incubated with polyclonal rabbit anti-BDNF (1:1000 in TBST + 5% skim milk) at 4°C for 2 h and then with horseradish peroxidase (HRP)-conjugated donkey anti-rabbit IgG (1:10000 in TBST + 5% skim milk) at 4°C for 1 h. The signals were detected using an enhanced chemoluminescence system (Amersham Biosciences, NJ) and visualized through X-ray film exposure (Kodak). The membranes were then stripped using Restore Western Blot Stripping Buffer (Pierce, Rockford, IL) for 4 - 6 hours at room temperature and re-probed with anti-β actin antibody (1:2000 in TBST + 5% skim milk) followed by HRP-conjugated donkey anti-goat IgG (1:15000 in TBST + 5% skim milk). The exposed X-ray films were digitized and densitometric quantification was performed using ImageJ (National Institute of Health, Bethesda, MD).

The BDNF levels were normalized to corresponding β actin signals, and the deduced ratios were further normalized to that of the control ND mouse on the same blot to perform statistical analyses with ANOVA followed by Bonferroni’s *post hoc* test. Rabbit polyclonal anti-BDNF (N-20 sc-546) and goat polyclonal anti-β actin (C-11 sc-1615) antibodies were purchased from Santa Cruz Biotechnology (Dallas, TX). Secondary antibodies (donkey anti-rabbit IgG and donkey anti-goat IgG, HRP-conjugate) were from Jackson ImmunoResearch (West Grove, PA). All other chemicals were purchased from Sigma (St. Louis, MO) unless otherwise noted.

## Acknowledgements

Supported by NIH grants R01EY002874 and R01EY025174. We thank members of the Stryker lab for a critical reading and T. Tran for technical assistance.

## References

Bozzi Y, Pizzorusso T, Cremisi F, Rossi FM, Barsacchi G, Maffei L. Monocular deprivation decreases the expression of messenger RNA for brain-derived neurotrophic factor in the rat visual cortex. Neuroscience. 1995 Dec;69(4):1133–44. doi: 10.1016/0306-4522(95)00321-9. PMID: 8848102.

Cabelli RJ, Hohn A, Shatz CJ. Inhibition of ocular dominance column formation by infusion of NT-4/5 or BDNF. Science. 1995 Mar 17;267(5204):1662–6. doi: 10.1126/science.7886458. PMID: 7886458.

Cabelli RJ, Shelton DL, Segal RA, Shatz CJ. Blockade of endogenous ligands of trkB inhibits formation of ocular dominance columns. Neuron. 1997 Jul;19(1):63–76. doi: 10.1016/s0896-6273(00)80348-7. PMID: 9247264.

Cang J, Kalatsky VA, Löwel S, Stryker MP. Optical imaging of the intrinsic signal as a measure of cortical plasticity in the mouse. Vis Neurosci. 2005 Sep-Oct;22(5):685–91. doi: 10.1017/S0952523805225178. PMID: 16332279; PMCID: PMC2553096.

Castrén E, Zafra F, Thoenen H, Lindholm D. Light regulates expression of brain-derived neurotrophic factor mRNA in rat visual cortex. Proc Natl Acad Sci U S A. 1992 Oct 15;89(20):9444–8. doi: 10.1073/pnas.89.20.9444. PMID: 1409655; PMCID: PMC50148.

Chen X, Ye H, Kuruvilla R, Ramanan N, Scangos KW, Zhang C, Johnson NM, England PM, Shokat KM, Ginty DD. A chemical-genetic approach to studying neurotrophin signaling. Neuron. 2005 Apr 7;46(1):13–21. doi: 10.1016/j.neuron.2005.03.009. PMID: 15820690.

Espinosa JS, Stryker MP. Development and plasticity of the primary visual cortex. Neuron. 2012 Jul 26;75(2):230–49. doi: 10.1016/j.neuron.2012.06.009. PMID: 22841309; PMCID: PMC3612584.

Faulkner SD, Vorobyov V, Sengpiel F. Visual cortical recovery from reverse occlusion depends on concordant binocular experience. J Neurophysiol. 2006 Mar;95(3):1718–26. doi: 10.1152/jn.00912.2005. Epub 2005 Dec 14. PMID: 16354732.

Gordon JA, Stryker MP. Experience-dependent plasticity of binocular responses in the primary visual cortex of the mouse. J Neurosci. 1996 May 15;16(10):3274–86. doi: 10.1523/JNEUROSCI.16-10-03274.1996. PMID: 8627365; PMCID: PMC6579137.

Gottmann K, Mittmann T, Lessmann V. BDNF signaling in the formation, maturation and plasticity of glutamatergic and GABAergic synapses. Exp Brain Res. 2009 Dec;199(3-4):203–34. doi: 10.1007/s00221-009-1994-z. Epub 2009 Sep 24. PMID: 19777221.

Harward SC, Hedrick NG, Hall CE, Parra-Bueno P, Milner TA, Pan E, Laviv T, Hempstead BL, Yasuda R, McNamara JO. Autocrine BDNF-TrkB signaling within a single dendritic spine. Nature. 2016 Oct 6;538(7623):99–103. doi: 10.1038/nature19766. Epub 2016 Sep 28. PMID: 27680698; PMCID: PMC5398094.

Hess RF, Thompson B. Amblyopia and the binocular approach to its therapy. Vision Res. 2015 Sep;114:4–16. doi: 10.1016/j.visres.2015.02.009. Epub 2015 Apr 20. PMID: 25906685.

Horch HW, Katz LC. BDNF release from single cells elicits local dendritic growth in nearby neurons. Nat Neurosci. 2002 Nov;5(11):1177–84. doi: 10.1038/nn927. PMID: 12368805.

Ichisaka S, Katoh-Semba R, Hata Y, Ohshima M, Kameyama K, Tsumoto T. Activity-dependent change in the protein level of brain-derived neurotrophic factor but no change in other neurotrophins in the visual cortex of young and adult ferrets. Neuroscience. 2003;117(2):361–71. doi: 10.1016/s0306-4522(02)00771-6. PMID: 12614676.

Jin L, Fang Y, Jin C. Binocular treatment for individuals with amblyopia: A systematic review and meta-analysis. Medicine (Baltimore). 2022 Jul 8;101(27):e28975. doi: 10.1097/MD.0000000000028975. PMID: 35801758; PMCID: PMC9259175.

Kaneko M, Hanover JL, England PM, Stryker MP. TrkB kinase is required for recovery, but not loss, of cortical responses following monocular deprivation. Nat Neurosci. 2008 Apr;11(4):497–504. doi: 10.1038/nn2068. Epub 2008 Mar 2. PMID: 18311133; PMCID: PMC2413329. (2008a).

Kaneko M, Stellwagen D, Malenka RC, Stryker MP. Tumor necrosis factor-alpha mediates one component of competitive, experience-dependent plasticity in developing visual cortex. Neuron. 2008 Jun 12;58(5):673–80. doi: 10.1016/j.neuron.2008.04.023. PMID: 18549780; PMCID: PMC2884387. (2008b).

Kaneko M, Cheetham CE, Lee YS, Silva AJ, Stryker MP, Fox K. Constitutively active H-ras accelerates multiple forms of plasticity in developing visual cortex. Proc Natl Acad Sci U S A. 2010 Nov 2;107(44):19026–31. doi: 10.1073/pnas.1013866107. Epub 2010 Oct 11. PMID: 20937865; PMCID: PMC2973899.

Kaneko M, Xie Y, An JJ, Stryker MP, Xu B. Dendritic BDNF synthesis is required for late-phase spine maturation and recovery of cortical responses following sensory deprivation. J Neurosci. 2012 Apr 4;32(14):4790–802. doi: 10.1523/JNEUROSCI.4462-11.2012. PMID: 22492034; PMCID: PMC3356781.

Kalatsky VA, Stryker MP. New paradigm for optical imaging: temporally encoded maps of intrinsic signal. Neuron. 2003 May 22;38(4):529–45. doi: 10.1016/s0896-6273(03)00286-1. PMID: 12765606.

Karpova NN, Rantamäki T, Di Lieto A, Lindemann L, Hoener MC, Castrén E. Darkness reduces BDNF expression in the visual cortex and induces repressive chromatin remodeling at the BDNF gene in both hippocampus and visual cortex. Cell Mol Neurobiol. 2010 Oct;30(7):1117–23. doi: 10.1007/s10571-010-9544-6. Epub 2010 Jul 8. PMID: 20614233.

Kind PC, Mitchell DE, Ahmed B, Blakemore C, Bonhoeffer T, Sengpiel F. Correlated binocular activity guides recovery from monocular deprivation. Nature. 2002 Mar 28;416(6879):430–3. doi: 10.1038/416430a. PMID: 11919632.

Kowiański P, Lietzau G, Czuba E, Waśkow M, Steliga A, Moryś J. BDNF: A Key Factor with Multipotent Impact on Brain Signaling and Synaptic Plasticity. Cell Mol Neurobiol. 2018 Apr;38(3):579–593. doi: 10.1007/s10571-017-0510-4. Epub 2017 Jun 16. PMID: 28623429; PMCID: PMC5835061.

Lein ES, Shatz CJ. Rapid regulation of brain-derived neurotrophic factor mRNA within eye-specific circuits during ocular dominance column formation. J Neurosci. 2000 Feb 15;20(4):1470–83. doi: 10.1523/JNEUROSCI.20-04-01470.2000. PMID: 10662837; PMCID: PMC6772351.

LeVay S, Wiesel TN, Hubel DH. The development of ocular dominance columns in normal and visually deprived monkeys. J Comp Neurol. 1980 May 1;191(1):1–51. doi: 10.1002/cne.901910102. PMID: 6772696.

Lessmann V, Gottmann K, Malcangio M. Neurotrophin secretion: current facts and future prospects. Prog Neurobiol. 2003 Apr;69(5):341–74. doi: 10.1016/s0301-0082(03)00019-4. Erratum in: Prog Neurobiol. 2004 Feb;72(2):165-6. PMID: 12787574.

Liu C, Chan CB, Ye K. 7,8-dihydroxyflavone, a small molecular TrkB agonist, is useful for treating various BDNF-implicated human disorders. Transl Neurodegener. 2016 Jan 6;5:2. doi: 10.1186/s40035-015-0048-7. PMID: 26740873; PMCID: PMC4702337.

Lu B. BDNF and activity-dependent synaptic modulation. Learn Mem. 2003 Mar-Apr;10(2):86–98. doi: 10.1101/lm.54603. PMID: 12663747; PMCID: PMC5479144.

Majdan M, Shatz CJ. Effects of visual experience on activity-dependent gene regulation in cortex. Nat Neurosci. 2006 May;9(5):650–9. doi: 10.1038/nn1674. Epub 2006 Apr 2. PMID: 16582906.

Mansouri B, Singh P, Globa A, Pearson P. Binocular training reduces amblyopic visual acuity impairment. Strabismus. 2014 Mar;22(1):1–6. doi: 10.3109/09273972.2013.877945. PMID: 24564723.

Meier K, Tarczy-Hornoch K. Recent Treatment Advances in Amblyopia. Annu Rev Vis Sci. 2022 Apr 4. doi: 10.1146/annurev-vision-100720-022550. Epub ahead of print. PMID: 35378045.

Meyer-Franke A, Kaplan MR, Pfrieger FW, Barres BA. Characterization of the signaling interactions that promote the survival and growth of developing retinal ganglion cells in culture. Neuron. 1995 Oct;15(4):805–19. doi: 10.1016/0896-6273(95)90172-8. PMID: 7576630.

Meyer-Franke A, Wilkinson GA, Kruttgen A, Hu M, Munro E, Hanson MG Jr, Reichardt LF, Barres BA. Depolarization and cAMP elevation rapidly recruit TrkB to the plasma membrane of CNS neurons. Neuron. 1998 Oct;21(4):681–93. doi: 10.1016/s0896-6273(00)80586-3. PMID: 9808456; PMCID: PMC2693071.

Mitchell DE, Gingras G, Kind PC. Initial recovery of vision after early monocular deprivation in kittens is faster when both eyes are open. Proc Natl Acad Sci U S A. 2001 Sep 25;98(20):11662–7. doi: 10.1073/pnas.201392698. PMID: 11573003; PMCID: PMC58786.

Mitchell DE, Duffy KR. The case from animal studies for balanced binocular treatment strategies for human amblyopia. Ophthalmic Physiol Opt. 2014 Mar;34(2):129–45. doi: 10.1111/opo.12122. PMID: 24588531.

Miyasaka Y, Yamamoto N. Neuronal Activity Patterns Regulate Brain-Derived Neurotrophic Factor Expression in Cortical Cells *via* Neuronal Circuits. Front Neurosci. 2021 Dec 10;15:699583. doi: 10.3389/fnins.2021.699583. PMID: 34955705; PMCID: PMC8702648.

Murphy KM, Roumeliotis G, Williams K, Beston BR, Jones DG. Binocular visual training to promote recovery from monocular deprivation. J Vis. 2015 Jan 8;15(1):15.1.2. doi: 10.1167/15.1.2. PMID: 25572348.

Pankiewicz P, Szybiński M, Kisielewska K, Gołębiowski F, Krzemiński P, Rutkowska-Włodarczyk I, Moszczyński-Pętkowski R, Gurba-Bryśkiewicz L, Delis M, Mulewski K, Smuga D, Dominowski J, Janusz A, Górka M, Abramski K, Napiórkowska A, Nowotny M, Dubiel K, Kalita K, Wieczorek M, Pieczykolan J, Matłoka M. Do Small Molecules Activate the TrkB Receptor in the Same Manner as BDNF? Limitations of Published TrkB Low Molecular Agonists and Screening for Novel TrkB Orthosteric Agonists. Pharmaceuticals (Basel). 2021 Jul 21;14(8):704. doi: 10.3390/ph14080704. PMID: 34451801; PMCID: PMC8398766.

Park H, Poo MM. Neurotrophin regulation of neural circuit development and function. Nat Rev Neurosci. 2013 Jan;14(1):7–23. doi: 10.1038/nrn3379. PMID: 23254191.

Pattabiraman PP, Tropea D, Chiaruttini C, Tongiorgi E, Cattaneo A, Domenici L. Neuronal activity regulates the developmental expression and subcellular localization of cortical BDNF mRNA isoforms in vivo. Mol Cell Neurosci. 2005 Mar;28(3):556–70. doi: 10.1016/j.mcn.2004.11.010. PMID: 15737745.

Reichardt LF. Neurotrophin-regulated signalling pathways. Philos Trans R Soc Lond B Biol Sci. 2006 Sep 29;361(1473):1545–64. doi: 10.1098/rstb.2006.1894. PMID: 16939974; PMCID: PMC1664664.

Repka, MX. Amblyopia: basics, questions, and practical management. Ch 73, pp. 754–761 in Taylor and Hoyt’s Pediatric Ophthalmology and Strabismus, 6th ed., Lambert SR, Lyons CJ (eds), Elsevier: Edinborough (2022)

Rossi FM, Bozzi Y, Pizzorusso T, Maffei L. Monocular deprivation decreases brain-derived neurotrophic factor immunoreactivity in the rat visual cortex. Neuroscience. 1999 May;90(2):363–8. doi: 10.1016/s0306-4522(98)00463-1. PMID: 10215141.

Schoups AA, Elliott RC, Friedman WJ, Black IB. NGF and BDNF are differentially modulated by visual experience in the developing geniculocortical pathway. Brain Res Dev Brain Res. 1995 May 26;86(1-2):326–34. doi: 10.1016/0165-3806(95)00043-d. PMID: 7656424.

Schwartz N, Schohl A, Ruthazer ES. Activity-dependent transcription of BDNF enhances visual acuity during development. Neuron. 2011 May 12;70(3):455–67. doi: 10.1016/j.neuron.2011.02.055. PMID: 21555072.

Specht KM, Shokat KM. The emerging power of chemical genetics. Curr Opin Cell Biol. 2002 Apr;14(2):155–9. doi: 10.1016/s0955-0674(02)00317-4. PMID: 11891113.

Stellwagen D, Malenka RC. Synaptic scaling mediated by glial TNF-alpha. Nature. 2006 Apr 20;440(7087):1054–9. doi: 10.1038/nature04671. Epub 2006 Mar 19. PMID: 16547515.

Sun YJ, Espinosa JS, Hoseini MS, Stryker MP. Experience-dependent structural plasticity at pre-and postsynaptic sites of layer 2/3 cells in developing visual cortex. Proc Natl Acad Sci U S A. 2019 Oct 22;116(43):21812–21820. doi: 10.1073/pnas.1914661116. Epub 2019 Oct 7. PMID: 31591211; PMCID: PMC6815154.

Tanaka J, Horiike Y, Matsuzaki M, Miyazaki T, Ellis-Davies GC, Kasai H. Protein synthesis and neurotrophin-dependent structural plasticity of single dendritic spines. Science. 2008 Mar 21;319(5870):1683–1687. doi: 10.1126/science.1152864. Epub 2008 Feb 28. Erratum in: Science. 2009 Dec 11;326(5959):1482. PMID: 18309046; PMCID: PMC4218863.

Tongiorgi E. Activity-dependent expression of brain-derived neurotrophic factor in dendrites: facts and open questions. Neurosci Res. 2008 Aug;61(4):335–46. doi: 10.1016/j.neures.2008.04.013. Epub 2008 May 4. PMID: 18550187.

Trachtenberg JT, Trepel C, Stryker MP. Rapid extragranular plasticity in the absence of thalamocortical plasticity in the developing primary visual cortex. Science. 2000 Mar 17;287(5460):2029–2032. doi: 10.1126/science.287.5460.2029. PMID: 10720332; PMCID: PMC2412909.

Tuszynski, M. A clinical trial of AAV2-BDNF gene therapy in early Alzheimer’s disease and mild cognitive impairment. https://www.clinicaltrials.gov/ct2/show/NCT05040217 (2022)

Wiesel TN. Postnatal development of the visual cortex and the influence of environment. Nature. 1982 Oct 14;299(5884):583–91. doi: 10.1038/299583a0. PMID: 6811951.

